# Predicting protein functional motions: an old recipe with a new twist

**DOI:** 10.1101/703652

**Authors:** Sergei Grudinin, Elodie Laine, Alexandre Hoffmann

## Abstract

Large macromolecules, including proteins and their complexes, very often adopt multiple conformations. Some of them can be seen experimentally, for example with X-ray crystallography or cryo-electron microscopy. This structural heterogeneity is not occasional and is frequently linked with specific biological function. Thus, the accurate description of macromolecular conformational transitions is crucial for understanding fundamental mechanisms of life’s machinery. We report on a real-time method to predict such transitions by extrapolating from instantaneous eigen-motions, computed using the normal mode analysis, to a series of twists. We demonstrate the applicability of our approach to the prediction of a wide range of motions, including large collective opening-closing transitions and conformational changes induced by partner binding. We also highlight particularly difficult cases of very small transitions between crystal and solution structures. Our method guaranties preservation of the protein structure during the transition and allows to access conformations that are unreachable with classical normal mode analysis. We provide practical solutions to describe localized motions with a few low-frequency modes and to relax some geometrical constraints along the predicted transitions. This work opens the way to the systematic description of protein motions, whatever their degree of collectivity. Our method is available as a part of the NOn-Linear rigid Block (NOLB) package at https://team.inria.fr/nano-d/software/nolb-normal-modes/.

**Significance Statement:** Proteins perform their biological functions by changing their shapes and interacting with each other. Getting access to these motions is challenging. In this work, we present a method that generates *plausible* physics-based protein motions and conformations. We model a protein as a network of atoms connected by springs and deform it along the least-energy directions. Our main contribution is to perform the deformations in a nonlinear way, through a series of twists. This allows us to produce a wide range of motions, some of them previously inaccessible, and to preserve the structure of the protein during the motion. We are able to simulate the opening or closing of a protein and the changes it undergoes to adapt to a partner.

---

Large macromolecules, including proteins and their complexes, are intrinsically flexible, and this flexibility is often linked with their function. A molecule in solution can be viewed as a structurally heterogeneous ensemble, where a finite number of conformational states (*e.g*. active-inactive, bound-unbound) may become stable under certain conditions to perform specific tasks. Identifying the molecular states relevant to protein functioning is necessary for our understanding of biological processes. Moreover, targeting protein functional motions bears a great potential to control and modulate proteins’ activities and interactions in physio-pathological contexts.

Structural heterogeneity can be probed by various experimental techniques. These include X-ray crystallography, cryo-electron microscopy (cryo-EM), nuclear magnetic resonance (NMR), small-angle scattering and many others (1). The two first methods allow obtaining large macromolecular structures at high resolution. While X-ray crystallography captures single stable states, cryo-EM allows observing conformational ensembles in solution. The resolution attained by cryo-EM is very often lower than that of X-ray structures, mainly due to the *structural heterogeneity* of the measured samples. However, the ongoing revolution in cryo-EM instrumentation (2) has supplied an exponentially growing body of near-atomic resolution structures. These techniques provide valuable insights on proteins’ functioning and interactions with their environment. Nevertheless, experimental protein structure determination remains a time consuming and costly process. The *systematic* description of the variety of shapes a protein adopts under particular environmental conditions, upon post-translational modifications and/or partner binding still remains out of reach. Hence, there is a need for computational tools able to efficiently and accurately predict functionally relevant protein conformations and macromolecular motions in general.

Several decades ago, Hayward and Go (3) observed that large-scale protein dynamics can be described with a set of just a few *collective coordinates*, accessible through the normal mode analysis (NMA). Thus, the latter provides an efficient way for reducing the dimensionality of the initial system and allows to study conformational transitions in proteins and their complexes. This has motivated the development of NMA-based tools for multiple biological applications, including flexible fitting of atomistic structures into cryo-EM maps (4–11) or one-dimensional scattering profiles (12), prediction of crystallographic temperature factors (13–15), generation of structural ensembles for cross-docking (16, 17), prediction of protein hinge regions (18, 19), flexible docking (20–23), refinement of crystallographic structures (24, 25) and docking solutions (26–28), and many others. The suitability of the NMA to model conformational dynamics varies widely depending on the system studied and on the type of motions involved (29). The NMA was shown to better describe highly collective motions, compared to localized deformations (30).

Atomistic molecular dynamics (MD) simulations represent an alternative to the NMA. They provide a practical tool to describe the structural heterogeneity around an equilibrium state and the flexibility exhibited by solvent-exposed small regions, such as loops. For instance, MD-based sampling has been applied to model the conformational diversity embedded in localized regions of cryo-EM maps (31). In addition, the concept of collective coordinates has been extended to MD (32–34), which, as a result, have been applied to the study of free energy changes between different conformational states, and rare-event dynamics (35). Nevertheless, MD simulations are much more costly than the NMA and the systematic characterization of conformational transitions with the former still remains computationally prohibitive.

In this work, we present an efficient real-time method to predict biomolecular transitions involving a wide range of motions, from local deformations, *e.g*. of a small loop, to highly collective domain motions. It relies on the nonlinear rigid block (NOLB) NMA (36). NOLB extends the classical NMA to describe nonlinear motions. Specifically, it extrapolates motions computed from instantaneous linear and angular velocities to large amplitudes. The resulting molecular motion is represented as a series of rigid block twists. We apply this nonlinear extrapolation to a combination of a few low-frequency normal modes to approximate conformational transitions. Importantly, our approach is conceptually simple and explores the conformational space in the Cartesian coordinate system. The nonlinearity of the computed motions allows a better approximation of experimentally observed transitions.

So far, the computation of nonlinear transitions using the NMA formalism has only been possible by cutting them in small steps and recomputing the normal modes at each step, and/or by performing the NMA in the internal coordinate system (11, 14, 37, 38). On average, the internal-coordinate NMA (iNMA) requires a smaller number of modes than the classical Cartesian-coordinate NMA to describe large structural transitions (14), and better predicts transitions upon protein docking (38). Working with internal coordinates also allows for large dimensionality reduction through variable selection and model simplification (14, 39–43). Despite these advantages, iNMA implies solving the generalized eigenvalue problem and dealing with necessarily dense interaction matrices. This makes it computationally costly and prevents its application on a large scale. Moreover, small changes in the internal coordinates may result in very large overall structural changes, which makes the approach less amenable to conformational space exploration, as it generates instability in the solution.

To demonstrate the advantages of the method reported here, we assess structural transitions computed with the classical linear normal modes, the Cartesian nonlinear normal modes, and an iterative scheme where the nonlinear modes are updated while progressing to the target state. For this purpose, we composed three test benchmarks of proteins exhibiting various types of structural transitions. The first test case presents examples of large domain motions, where ‘open’ and ‘closed’ conformations can be clearly identified (11). The second one is comprised of proteins changing their conformation upon binding to other proteins (44). The third one contains test cases from the Cryo-EM 2015/2016 Model Challenge, where the transition takes place between a crystal form and a conformation in solution (45). We find that the classical linear NMA behaves well on the first set, where the motions are mostly collective, but is not suited to describe the more localized deformations and very small transitions exhibited by the two other sets. We show that our Cartesian nonlinear approach systematically obtains better transitions compared to the linear one. Indeed, the final predicted structures are closer to the experimentally known targets and display less distortions. The improvement is particularly significant on changes associated to partner binding. We further demonstrate the usefulness of nonlinearity and mode updating to extend the applicability of the NMA to localized and disruptive motions. Last, but not least, our approach is very computationally and memory efficient. It is implemented as a fully automated tool available at: https://team.inria.fr/nano-d/software/nolb-normal-modes/.

Our results allow revisiting the NMA-based description of biomolecular transitions. They pave the way to the systematic targeting and modulation of protein-protein interactions.

## Computational method

Protein shapes and motions are governed by a multitude of interatomic forces, resulting from intra- and inter-molecular interactions. Despite this high complexity, many functional motions can be approximated by a few *low-frequency modes* characteristic of the protein’s geometrical shape (30, 46, 47). To compute these modes, we represent the protein as an elastic network (Fig. 1, top panel on the left), where each node stands for an atom and two nodes *i* and *j* are connected by a spring whenever the distance *d_i_j* between the corresponding atoms is smaller than a cutoff value, typically 5 Å (*SI Appendix, Text 2F*). The normal modes are obtained by diagonalizing the mass-weighted Hessian matrix of the potential energy of this network (*SI Appendix, Text 2A*). To reduce the dimensionality of this diagonalization problem, we consider each protein residue as a rigid block, according to the *rotation translation blocks* (RTB) approach (48, 49) (Fig. 1, middle panel on the left, see also *SI Appendix, Text 2B*). With this coarse-grained representation, the computed normal modes are composed of *instantaneous linear velocities* 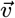 and *instantaneous angular velocities* 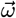, defining translations and rotations for each block/residue.

**Fig. 1.**
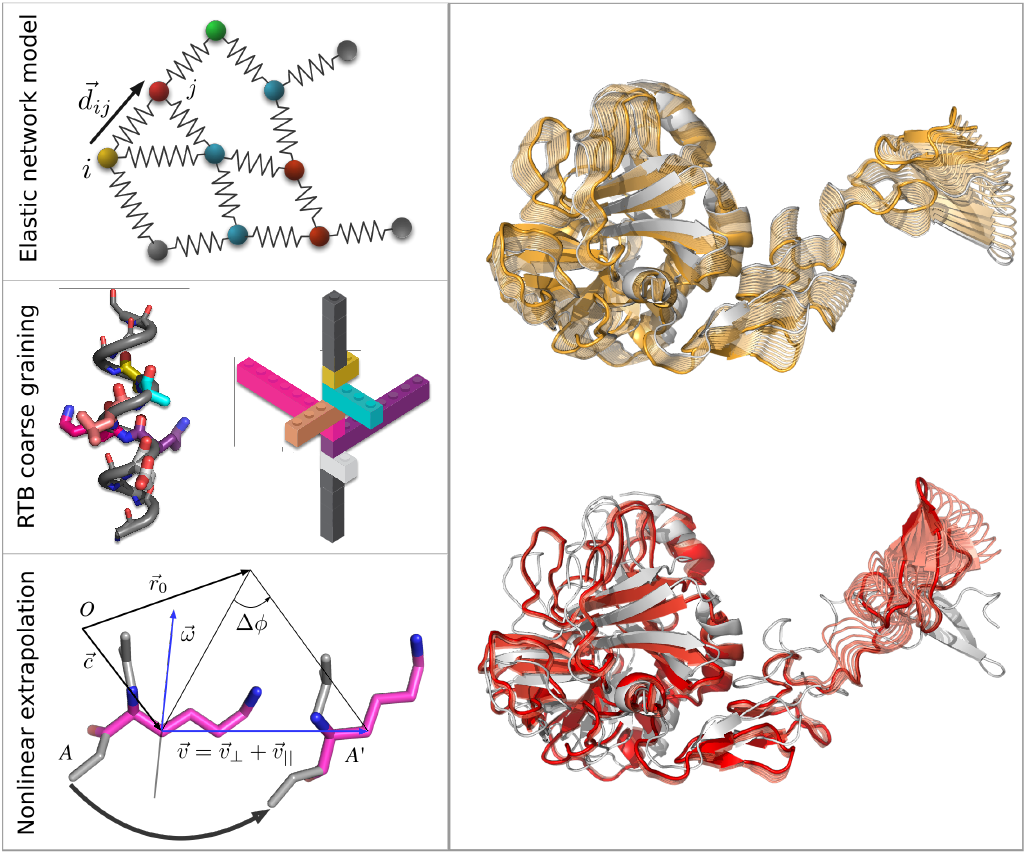
Principle of the method. Left panel. The three main ingredients of the method are depicted: the elastic network model, the rotation translation blocks (RTB) projection and the nonlinear extrapolation of motions. The protein is represented as an elastic network (on top), where all the atom pairs within a certain cutoff distance are connected with harmonic springs. Coarse-graining is achieved by replacing each protein residue by a rigid block (in the middle). The color code indicates the one-to-one correspondence between residues (on the left) and blocks (on the right). Each block has six degrees of freedom, three for rotation and three for translation. At each step of the transition, each residue/block undergoes a screw (twist) motion (at the bottom) defined from the instantaneous linear and angular velocities 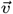 and 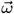 obtained by the NMA. The initial and final atomic positions are denoted as *A* and *A*′, respectively. *O* is the origin of the coordinate system and c is the residue’s center of mass. **Right panel.** Examples of nonlinear transitions computed for coagulation factor VIIa upon binding to tissue factor. The intermediate structures in orange were determined from the normal modes of the known unbound structure (1qfk:HL, in grey). Those in red were further obtained by updating the normal modes three times. The final predicted structure (in opaque) is 1.3 Å from the known bound structure (1fak:HL).

A straightforward way to compute normal-mode guided structural transitions is to calculate instantaneous displacements of each atom in a residue and then linearly extrapolate these up to a given amplitude ***a***. However, at large amplitudes, this will distort interatomic distances and produce unrealistic molecular conformations. To circumvent this problem, we apply a nonlinear extrapolation (Fig. 1, bottom panel on the left), where each residue undergoes a *screw* (or a *twist)* motion (*SI Appendix, Text 2C*). Specifically, the linear velocity 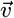 is decomposed in two terms, namely 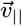, which is collinear to 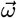, and 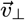, which is orthogonal to 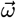. We further represent the pair of 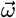 and 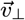 as a pure rotation around a new center 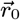. Hence, instead of rotating about the axis defined by 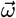 passing through its center of mass, each residue is rotated about the new axis defined by 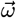 passing through 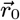 and translated only in the direction of 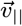 (*SI Appendix, Eq. 10*). This nonlinear extrapolation guarantees preservation of the topology or the protein structure subject to the motion.

Our method computes normal mode-guided nonlinear conformational transitions, starting from an experimentally determined structure or a high-quality 3D model. Specifically, normal modes are computed from the starting structure, which is then deformed along a selection of these modes up to a given amount of conformational deviation (Fig. 1, right panel, see also *SI Appendix, Text 2E*). The simulated conformational change can be potentially very large (several tens of Å). The algorithm may be run in an iterative mode, where the normal modes are re-computed on intermediate conformations. This allows modifying the topology of the network representing the structure and going further away from the starting structure (Fig. 1, right panel, compare orange and red conformations). The method guarantees producing *plausible* physics-based motions and conformations.

## Results

### NOLB nonlinear transitions improve the coverage of a wide range of functional motions

We assessed the nonlinear transitions computed by NOLB against 132 pairs of experimentally determined structures displaying a wide range of biologically relevant conformational changes. The root mean square deviation (RMSD) between the two structures range from 0.5 Å to 33 Å and the motions involve up to 80% of the protein atoms. For each pair, we defined a starting structure and a target structure. For a subset of 23 pairs (open-closed set, see below), each structure alternatively played the role of the starting structure and the target structure, resulting in a total of 155 predicted transitions. The transitions were systematically computed by using the ten lowest-frequency modes from the starting structure. In the general case, the target is not known and one has to sample the amplitudes of the modes. Here, we place ourself in a context where the amplitudes are determined by using the knowledge of the displacement between the starting and target structures (*SI Appendix, Eq. 13*). This allows obtaining the optimal (or close-to-optimal) transitions within our framework. We should mention that the method is very rapid. To compute all transitions reported here, it took us less than 5 minutes with one iteration, and about 15 minutes with five iterations, on a single CPU (*SI Appendix, Text 4*). The quality of a prediction was evaluated by computing its *transition coverage, i.e*. the relative RMSD explained by the prediction (*SI Appendix, Text 3A*). For instance, given a transition of 5 Å, a prediction achieving a coverage of 70% will produce a final conformation 1.5 Å away from the target structure.

On average, the NOLB nonlinear normal modes, computed with five iterations, covered 48% of the transitions. For comparison, the average coverage obtained with the classical linear modes was 40%. Moreover, the nonlinear predictions better approximated the transitions in 92% of the cases (Fig. 2A). The superiority of the NOLB predictions was also found significant without any update of the modes along the transition (*SI Appendix, Figure S1*). The anticoagulation factor VIIa (Fig. 1, right panel, and *Movie S1*) gives an illustrative example of a large transition (6.2 Å) upon binding to a cellular partner. The transition involves a complex motion of an “arm” comprising about 20% of the protein. The classical linear modes covered one third of the transition, producing a conformation 4.1 Å away from the target. The nonlinear NOLB normal modes achieved 44% coverage (Fig. 1, conformations in orange) and 79% after updating the modes 3 times (conformations in red). The final conformation is only 1.3 Å away from the target.

**Fig. 2.**
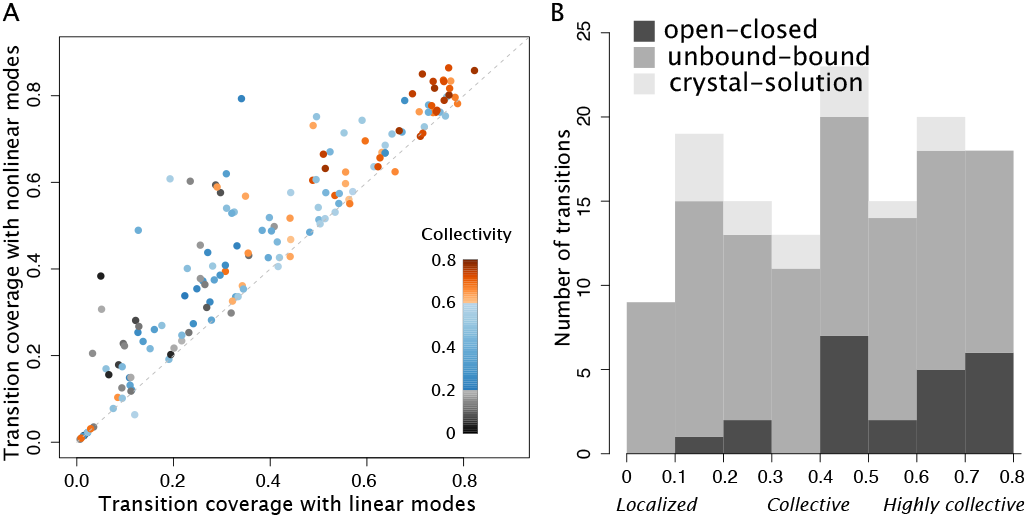
Transition coverage and collectivity. **A.** Comparison of the coverage achieved by the NOLB nonlinear modes with 5 iterations (y-axis) versus the classical linear modes (x-axis) for 155 transitions corresponding to 132 structure pairs (both forward and reverse transitions were computed for a subset of 23 pairs). The colors indicate the degrees of collectivity of the experimental transitions. **B.** Histogram of the collectivity degrees for all structure pairs from the three test sets. The transitions are labelled as localized (below 0.2), collective (between 0.2 and 0.6) and highly collective (above 0.6).

### NOLB extends the applicability of the normal mode analysis to localized motions

We collected the pairs of experimental structures from three benchmark sets (*SI Appendix, Text 1*) designed for different practical applications, namely NMA, docking and cryo-EM fitting. The first set comprises 23 proteins undergoing opening/closing motions. The vast majority of these transitions involve more than 40% of the protein atoms (Fig. 2B, dark grey bars). They can be explained by a few low-frequency normal modes (typically 1-3) computed from the open form (Fig. 3A, see bars in blue tones on the left). The second set contains 95 structural transitions associated to the binding of a protein partner. Such transitions are particularly challenging for protein docking applications (20, 21, 23, 50–52). Indeed, they are often induced by the spatial proximity of the partner (induced-fit mechanism), which makes them very difficult to estimate starting only from the knowledge of the unbound state. This set includes a great variety of motions, from highly localized to highly collective ones (Fig. 2B, medium grey bars). The transition coverage achieved by the classical linear normal modes is rather low (below 40%) for the majority of transitions (Fig. 3B, see colored bars). The few transitions explained by the first three modes (see right part of the plot) involve more than 70% of the protein atoms and are all antibodies. The third set comprises 14 transitions between either a crystal structure and a solution structure solved by cryo-EM or between two cryo-EM solution structures (*SI Appendix, Table S1*). Contrary to the other two sets, it is dominated by very small transitions (<2 Å, see *SI Appendix, Fig. S2*). The explanative power of the 10 first modes is very poor on this set (Fig. 3C).

**Fig. 3.**
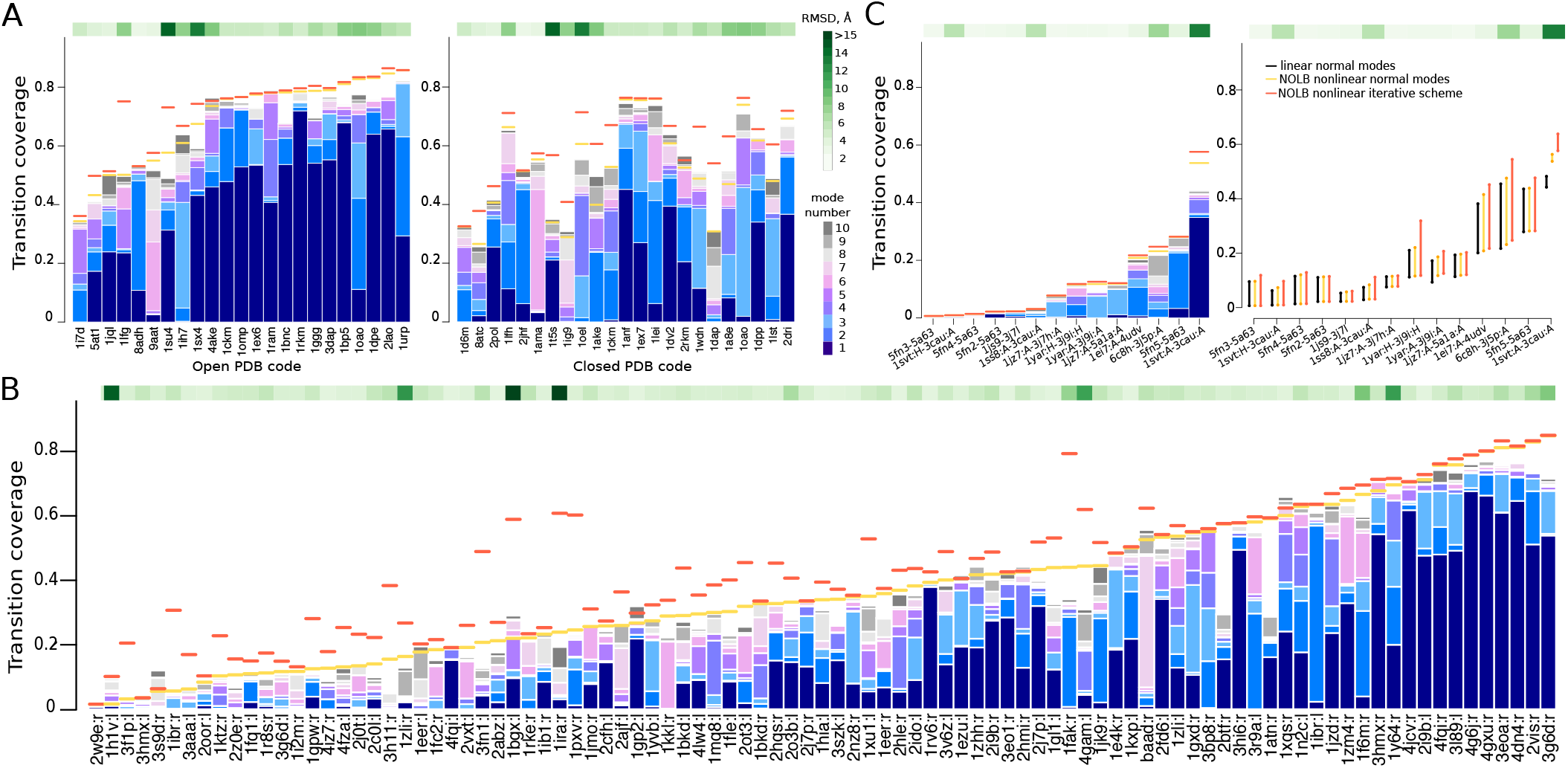
Comparison of the transition coverages achieved on the three benchmark sets. The strips on top show *C_α_* RMSD between the two known structures. The *y*-axes show the transition coverage achieved by the 10 lowest-frequency linear normal modes (bars in blue tones), the NOLB nonlinear normal modes (in orange) and the NOLB nonlinear iterative scheme (in red). The *x*-axes list the PDB codes, ordered according the NOLB normal mode predictions’ quality **A.** Open-to-closed (on the left) and closed-to-open (on the right) transitions. **B.** Unbound-to-bound transitions. **C.** Crystal-to-solution transitions. The plot on the right shows the improvement of the predictions when increasing the number of active normal modes from 10 to 40.

Overall, the ability of the classical NMA to predict transitions is largely determined by the transitions’ collectivity degrees (Fig. 2A, see the color gradient along the x-axis). Highly collective motions tend to be very well predicted while localized motions tend to be poorly predicted, in agreement with previous works (14, 30, 53). We found that our nonlinear scheme permits to *go beyond this observation and extends the applicability of the NMA*. Indeed, the highest improvement of NOLB predictions over the classical NMA is observed for localized transitions, involving less than 20% of the protein atoms (Fig. 2A, grey dots). The transition coverage is more than twice as big, on average, reaching a maximum value of 60% (versus 40% for the linear normal modes). As illustrative examples, let us mention Ephrin B4 receptor (2hle:r), Cystein protease (1pxv:r), actin (1atn:r and 2btf:r) and Rabex-5 VPS9 domain (2ot3:l), which undergo localized motions upon binding to their partners (Fig. 3B, see the location of the orange and red segments). While the linear modes predict between 23 and 36% of their transitions, our nonlinear scheme predicts between 43 and 60% of them. The linear and nonlinear transitions predicted for actin are illustrated in Fig. 4A (in blue and orange, see also *Movies S2* and *S3*).

**Fig. 4.**
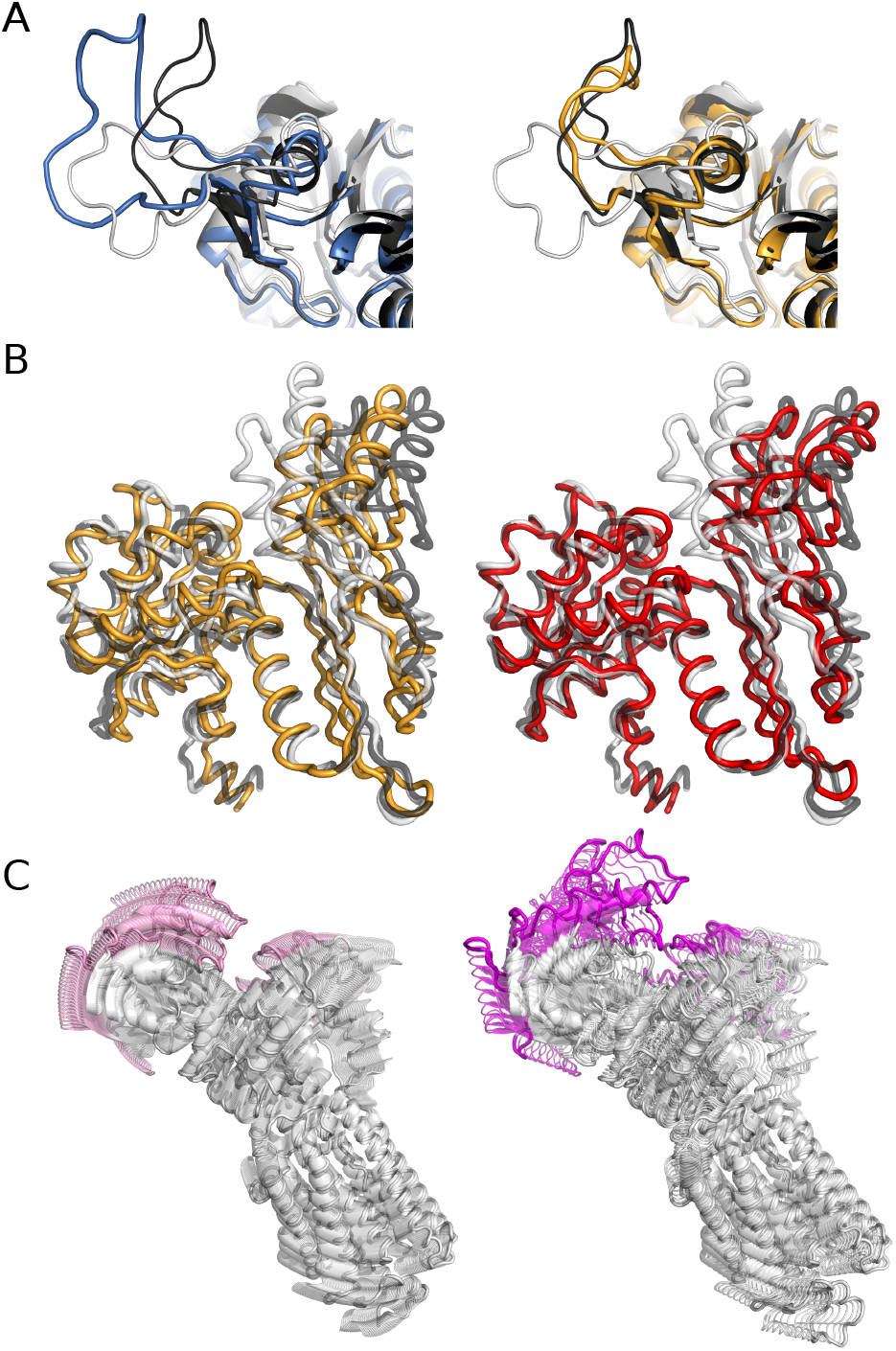
Examples of predicted transitions. **A.** Transition of actin upon binding to DNase I (1atn:r). The RMSD between the starting structure (in grey) and the target structure (in black) is 2.7 Å and the motion involves 10% of the protein atoms. The final conformation predicted by the classical NMA is displayed on the left in marine. The conformation predicted by NOLB nonlinear normal modes (without any update) is displayed on the right in orange. They deviate by 1.9 Å and 1.1 Å from the target, respectively. **B.** Opening of the diaminopimelate dehydrogenase (1dap–3dap). RMSD between the starting structure (in grey) and the target structure (in black) is 4.2 Å and the motion involves about half of the protein atoms. The final conformations predicted by NOLB are displayed in orange (without any mode updating, on the left) and in red (after one update, on the right). They deviate by 2.9 Å and 1.9 Å from the target structure, respectively. **C.** Closing of the calcium ATPase pump (1su4–1t5s). Conformations predicted by the NOLB nonlinear modes (without any update) and the classical NMA are shown on the left and the right, respectively. The residues undergoing the highest displacements are highlighted in color.

### Updating of the modes allows relaxing the elastic network’s constraints

The transitions predicted by the classical NMA strongly depend on the geometrical shape of the starting structure. This is particularly visible on the first test set, where the closed-to-open transitions are significantly worse than the open-to-closed ones (compare the two plots in Fig. 3A). Moreover, the number of transitions explained (at more than 40%) by the first three modes reduces from 18 to 8 upon starting from the closed structure. This effect was observed previously (30) and has a clear physical explanation connected to the limitations of the elastic network model. Indeed, in the closed state, this model contains a larger number of elastic links compared to the open state. Therefore, it is more difficult to produce a large deformation along a few directions from this more constrained starting point.

By re-computing the modes along the transition, our iterative scheme permits to overcome this limitation. Namely, it increases the coverage in the closed-to-open direction from 53% to 61%, on average (Fig. 3A, see the location of the red segments on the right). This result can be explained by the fact that, at each iteration, some elastic links are removed, alleviating some of the constraints that exert on the closed structure. As a consequence, the discrepancy between open-to-closed and closed-to-open predictions is largely reduced (compare the left and right plots). In four cases, namely the aspartate amino transferase (9aat-1ama), the maltodextrin binding protein (1omp–1anf), the alcohol dehydrogenase (8adh–2jhf) and the guanylate kinase (1ex6–1ex7), the coverage achieved in the two directions even becomes equivalent. The highest increase in coverage is obtained for the diaminopimelate dehydrogenase (1dap–3dap), from 31% without any update to 54% after one update (Fig. 4B, compare orange and red conformations, and see *Movie S4*).

### Nonlinear transitions better preserve the protein structure

Beyond improving the coverage of the transitions, the NOLB method produces motions that better preserve the overall protein structure and local topology. The predicted transitions visually look better than those produced with the linear extrapolation. The difference is particularly visible when dealing with large displacements. For instance, the calcium ATPase pump (1su4-1t5s) undergoes a large domain motion taking place during active transport. The RMSD between the open and closed conformations is of 13.5 Å. The nonlinear transition computed by NOLB, without any update of the modes, reaches a coverage of 58% while preserving very well the structure of the protein (Fig. 4C, on the left, and see *Movie S5*). Conversely, the linear transition attains only 49% coverage and it visibly distorts the cytoplasmic headpiece, where the closing motion takes place (Fig. 4C, on the right, and see *Movie S6*).

### Very small transitions remain difficult to predict

The third benchmark set, representing transitions between crystallographic structures and structures found in solution, was particularly challenging for the classical NMA and the NOLB method (Fig. 3C). On average, the ten first modes contribute to only 12% of the structural transitions. The improvement brought by the NOLB predictions is very limited. Using 40 active modes significantly improves the coverage, up to 26% of the transition. Nevertheless, this percentage still seems very low compared to the previous test cases and also for any practical applications. Most of the conformational changes from this set are of very small amplitude, even below 1 Å (*SI Appendix, Fig. S2*). This may explain the low coverages we obtain. The average RMSD between the final conformation computed by NOLB and the target structure is 1.9 Å. The only very large transition of the set, namely that of the GroEL chaperone (1svt:A–3cau:A), with an initial RMSD of 12.1 Å, is predicted at 58% by the NOLB iterative scheme.

## Conclusions

This work revisits the formalism of normal modes and demonstrates its applicability to the previously inaccessible cases of localized motions. Specifically, it critically assesses the relevance of the normal mode analysis to the computation of various structural transitions in biological macromolecules. Our results challenge the long-standing belief that the lowest-frequency modes can only describe collective transitions. Indeed, we show that nonlinear normal modes can also approximate local deformations such as loop motions. Moreover, iterative recomputation of the normal modes relaxes constraints imposed by the geometry of the protein and allows pushing the transitions even further. Another important advantage of our method is that the predicted conformations have a much better local geometry than those resulting from linear NMA perturbations.

Small structural changes, for example those present in the Cryo-EM 2015/2016 Model Challenge benchmark, still remain very difficult to predict with the NMA formalism. Indeed, in this case adding nonlinearity and iterative computations did not improve the results significantly. Activating a much larger number of modes can help approximating the transitions, but at the expense of a significant computational cost. Indeed, the full diagonalization of the Hessian matrix scales as *O*(*N*^3^) with the number of degrees of freedom *N*. Therefore, it becomes preferable to use MD-based or other stochastic optimization techniques, *i.e*. simulated annealing, with the full range of degrees of freedom.

Our method is very CPU and memory efficient – it took us about 9 minutes to compute the nonlinear structural transitions for all proteins from the PPDBv5 (460 in total) set on a desktop computer. This implies that the method can be applied on a very large scale. For instance, it can be used to model flexibility in docking calculations or to generate putative conformations that can be targeted by small molecules.

## Supporting Information (SI)

SI contains Figs. S1 to S2, Table S1, captions for Movies S1 to S6, and references for SI citations.

## SI Movies

Supporting movies S1-S6 can be found online.

## Materials and Methods

*SI Appendix, Text* includes detailed descriptions of the datasets (*SI Appendix, Text 1*), the computational framework (*SI Appendix, Text 2*), the assessment of the transitions (*SI Appendix, Text 3*) and the computational details, including command lines used to generate the transitions (*SI Appendix, Text 4*). The method is freely available as a part of the NOn-Linear rigid Block (NOLB) package at https://team.inria.fr/nano-d/software/nolb-normal-modes/. Scripts used to produce the reported data are also available at this address.

## Supporting Information Text

### 1. Datasets

#### Test set 1

For the first test set we used protein structures from the iMod benchmark (1) prepared by Chacón and colleagues (available at http://chaconlab.org/multiscale-simulations/imod/imod-donwload/item/imod-benchmark). It was recently used to assess three coarse-grained elastic network model-based flexible fitting methods (2). It comprises 23 proteins, each given in “open” and “closed” conformations, and represents a wide variety of macromolecular motions. While hinge motions are largely represented, the dataset also comprises shear and other complex motions. The structures were extracted from the molecular motions database MolMovDB (3). All of them have less than 3% Ramachandran outliers (as computed by the MolProbity program (4)), do not have any broken chain or missing atom. The average displacement for this test set is 5.1 ± 3.0 Å.

#### Test set 2

For the second test set we have chosen some examples from the Protein-Protein Docking Benchmark v5 (PPDBv5) (5). This benchmark contains 230 protein complexes with at least one of the partners solved in both bound (complexed) and unbound (free) states. All structures have a resolution better than 3.25 Å, and some of them contain more than one chain. We extracted 95 proteins with C_*α*_ RMSD displacements between the two states above 2 Å. This test set is well suited for assessing the range of applicability of flexible docking methods (6). We should also mention that some of the structure pairs can be classified as open-closed pairs. The average displacement for this test set is 4.0 ± 3.9 Å.

#### Test set 3

For the third test set we have selected seven cases from the Cryo-EM 2015/2016 Model Challenge (7). The initial set was comprised of eight cases, but we decided not to consider one of them, namely the 70S ribosome. The selected cases are listed in Table S1. Each one of them comprises one or several starting structures solved by X-ray crystallography and one or several target structures corresponding to a Model Challenge map. In one case (*γ*-secretase) we did not find homologous X-ray structures for the starting state and used several cryo-EM structures instead. The map resolutions range from 2.2 to 4.3 Å. The average C_*α*_ RMSD displacement between the two states is 2.6 ± 3.2 Å.

### 2. Computational model and framework

#### A. NMA theory

Let us consider a molecular system with ***N*** atoms at an equilibrium position ***q***_0_ ∈ ℝ^3***N***^. Let ***V***: ℝ^3***N***^ ↦ ℝ be the potential energy of the molecular system. Let us also introduce ***q*** ∈ ℝ^3***N***^, a small time-dependent displacement of the system around ***q***_0_. The potential energy ***V*** in the vicinity of ***q***_0_ can thus be given by its quadratic approximation, which allows to *analytically* solve the Newton’s equation of motion,

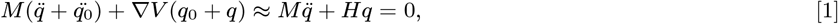

where ***M*** is the diagonal mass matrix, and ***H*** is the *Hessian matrix* of the potential energy ***V*** evaluated at the equilibrium position ***q***_0_. We then compute the square matrix of eigenvectors ***L*** and the diagonal matrix of eigenvalues Λ of the *mass-weighted* Hessian ***H***_*w*_ = *M*^−1/2^*HM*^−1/2^,

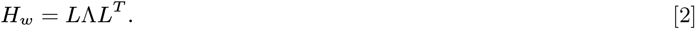

Let us now introduce ***η*** ∈ ℝ^3***N***^, a projection of ***q*** into the eigenspace of ***H***_*w*_, and (λ_*i*_)_***i***=0…3***N***_, the diagonal values in λ. Then, left multiplying Eq. 1 by ***L***^*T*^*M*^1/2^ gives the following system of uncoupled equations,

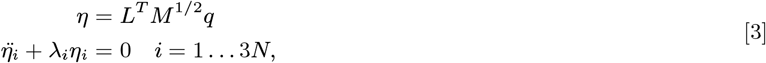

which can be solved analytically. We will refer to the columns of the ***M***^−1/2^***L*** matrix as to *Cartesian linear normal modes*. We should specifically mention that these normal modes are not generally orthogonal, unless all the masses in ***M*** are equal to each other.

#### B. The RTB projection method

Many methods have been proposed to reduce the dimensionality of the NMA diagonalization problem. For example, Noguti and Gõ (8) and Levitt et al. (9), and later Ma et al. (10), Mendez and Bastolla (11), and Chacón et al. (1) explored the NMA approach in internal coordinates. However, an orthogonal idea of reducing the dimensionality of the original system by coarse-graining its representation has gained much more popularity. One of the first coarse-graining methods was the *rotation translation blocks* (RTB) approach introduced by Durand et al. (12) and further developed by Tama et al. (13) and Li and Cui (14). In this method, individual or several consecutive amino residues are considered as rigid blocks that can only exhibit rotational and translational motions (12, 13). The transition from the RTB coordinate system, consisting of ***n*** rigid blocks with 6n DOFs to the all-atom coordinate system with 3N DOFs is performed by an *orthogonal projection matrix* ***P*** ∈ ℝ^3***N***×6***n***^, whose detailed form can be found elsewhere (15). We will only mention that this projection matrix is obtained by writing down the conservation laws of the linear and the angular momenta for a rigid block in mass-weighted coordinates (12).

The normal modes are then computed by the diagonalization of the RTB-projected mass-weighted Hessian,

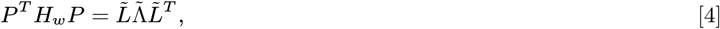

where 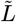 is the matrix composed of the RTB normal modes with the corresponding diagonal eigenvalue matrix 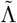. The all-atom normal modes ***L^w^*** (in mass-weighted coordinates) are then obtained as a projection of the RTB normal modes 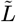 according to

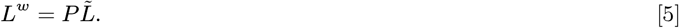

#### C. The nonlinear NOLB NMA method

Molecular vibrations in a multi-dimensional harmonic oscillator are all uncoupled and can be found by solving Eq. 3. Diagonalization of the RTB-projected mass-weighted Hessian gives a set of eigenvectors that are composed of *instantaneous linear velocities* 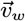 and *instantaneous angular velocities* 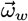 of individual rigid blocks. For a rigid block with mass ***M_b_*** and inertia tensor ***I***, we first compute these in non-mass weighted coordinates as follows,

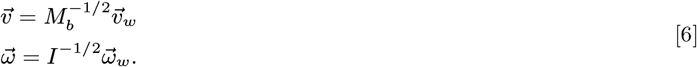

Then, given a deformation amplitude ***a***, the translational increment in the rigid block’s position 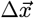 and the angular increment in its orientation Δ*ϕ* can be computed as

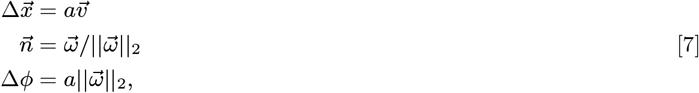

where the rigid block’s rotation is described with a unit axis 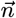 passing though its center of mass (COM) 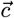, and an angle Finally, we rewrite the increment in the rigid block’s position 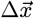 as a sum of two orthogonal vectors,

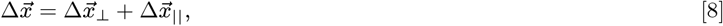

where 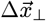 is orthogonal to 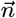, and 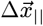 is collinear to 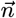. We then represent the 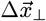-related motion as a pure rotation about a new center 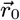 given as

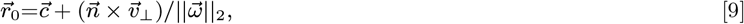

such that the final rigid block’s positions 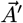 is expressed through the initial positions 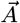 as

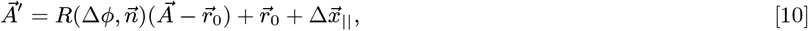

where 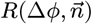 is the rotation matrix describing rigid block’s rotation about an axis 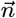 by an angle Δ*ϕ*. More details can be found in the original NOLB publication (15). It is easy to demonstrate that this is the only type of rigid-body motion that conserves the original kinetic energy. Indeed, using the parallel axis theorem it is readily seen that the initial energy contribution of linear velocity 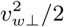 is transformed into equivalent contribution from the angular velocity. As it has been noted by Juan Cortés from LAAS-CNRS in a private communication, the presented theory can be also formulated in terms of *screw algebra*, where a screw is a six-dimensional vector constructed from a pair of three-dimensional vectors, linear and angular velocities.

#### D. Linear structural transitions

Let us assume that we have two conformations of the same molecular system with the known correspondence between the atoms in the two conformations. The correspondence can be robustly deduced using, *e.g*., sequence alignment of the two systems, if they are composed of not fully identical proteins. Let us also assume that we are given the displacement vector 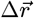 between the two conformations after their optimal rigid superposition. It is easy to demonstrate that in this case, the COMs of the two conformations match. We can now find the minimum root-mean-square deviation (RMSD) between the two conformations, if one of them is allowed to deform along its ***M*** lowest normal modes ***L*** ∈ ℝ^3***N***×***M***^, which are not necessarily orthonormal, as

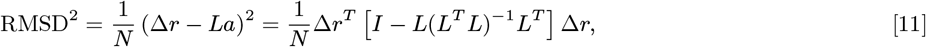

where ***N*** is the number of atoms in the system, ***I*** is the identity matrix, and ***a*** are the optimal amplitudes of linear deformations given as

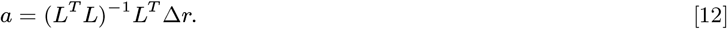

If the normal modes ***L*** are orthonormal (which happens if the mass matrix in Eq. 3 is identity), the above equation simplifies to

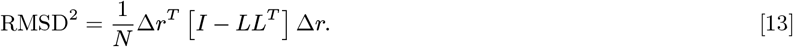

It can be readily seen that if all the 3***N*** modes are activated, the matrix ***L*** becomes square, ***LL^T^*** turns into an identity, and the RMSD reduces to zero.

#### E. Nonlinear structural transitions

The NOLB NMA method produces nonlinear deformations. Therefore, the RMSD equation 11 presented above would not be exact in this case. However, given the displacement vector 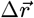 between the two conformations as in the previous case, we can still construct a deterministic deformation trajectory and compute the corresponding RMSD. We should specifically mention that rotation operators do not commute, and thus the result of application of two rotations would generally depend on the order of these operators. Therefore, to make the method deterministic, when combining multiple nonlinear motions corresponding to different normal modes, we have chosen to always apply the slower modes first. This choice is dictated by the fact that slower modes result in larger amplitudes of thermal fluctuations. Algorithm 1 lists steps producing a nonlinear deformation towards the target structure. In this algorithm, we use an iterative procedure, and at each step of the iteration we approximate the amplitudes of the nonlinear deformation by the analytically computed linear amplitudes using Eq. 12. This approximation will not be valid at large deformation amplitudes ***a***. Therefore, if the RMSD computed for the linear approximation (Eq. 11) is larger than a certain threshold (we have chosen 0.1 Å), we split the deformation into smaller pieces. Each piece is computed based on the values of the linear amplitudes scaled in such a way that the total linear RMSD of the deformation equals to the threshold value of 0.1 Å. We terminate the algorithm when the maximum number of iterations is exceeded (100 by default), or if the relative deformation becomes smaller than a tolerance of 1*e* − 6.

The abovementioned algorithm can be iterated multiple times. At each iteration, the elastic network model is updated and the normal modes are recomputed, as described in Algorithm 2. On-the-fly normal mode re-computation has been previously proposed in the context of cryo-EM fitting and morphing applications (1, 16, 17). We should specifically note that our nonlinear model and the way we assess the predicted transitions naturally overcome the limitations of classical NMA schemes highlighted in Jernigan et al (18, 19) when the transition involves a substantial protein domain rotation.

##### Algorithm 1. This deterministic algorithm produces a nonlinear structural deformation towards the target conformation

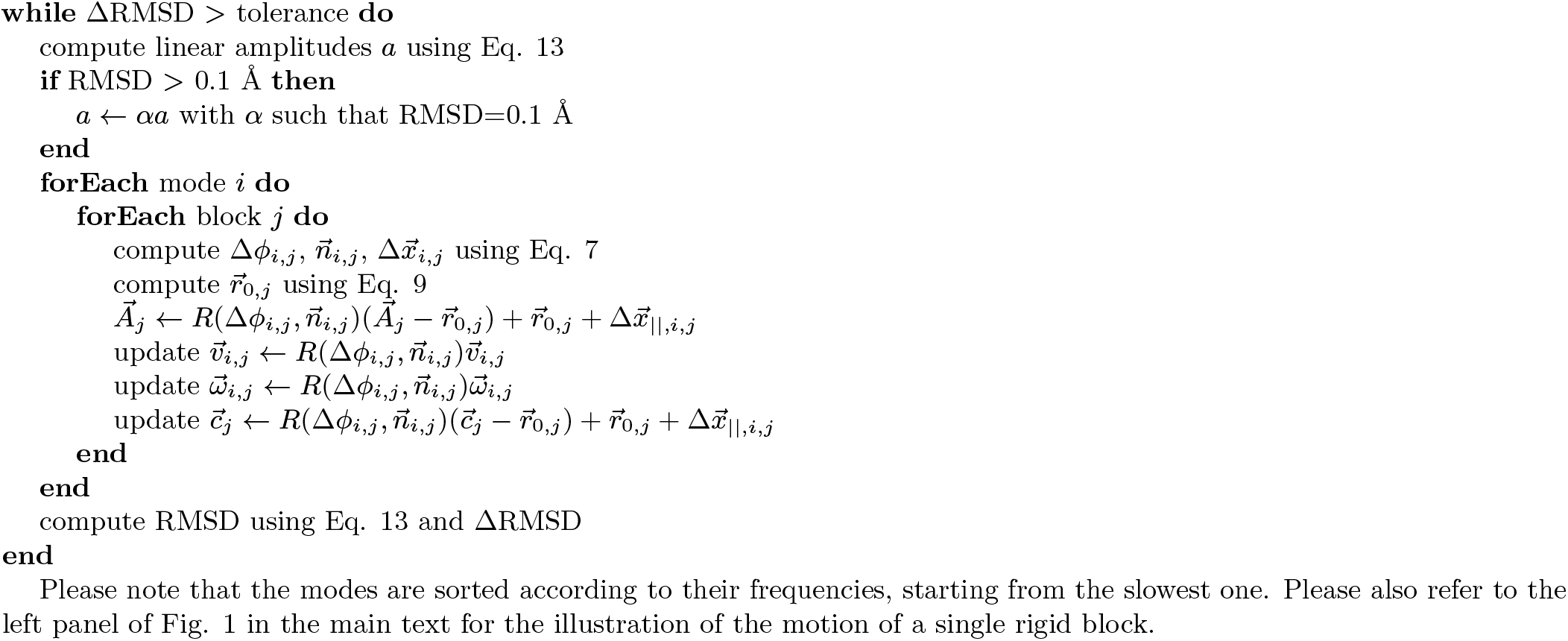

##### Algorithm 2. This extension of the previous algorithm produces a nonlinear structural deformation with multiple updates of the Hessian matrix

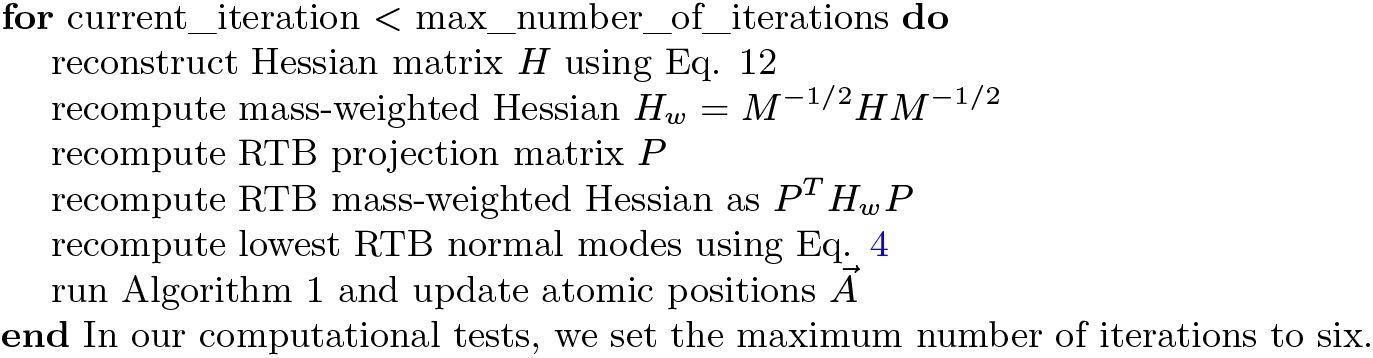

#### F. Potential function

Classical NMA methods can use any potential function, provided that it corresponds to the equilibrium position of the molecular system. Some recent developments can also assume non-equilibrium state of the initial system (20). In our method we use an all-atom anisotropic network model (ANM) (21, 22), where the initial structure is always at equilibrium. The all-atom ANM has the following potential function,

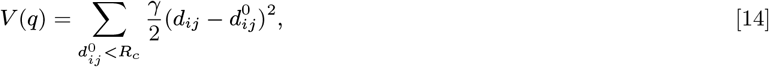

where ***d_ij_*** is the distance between the ***i^th^*** and the ***j^th^*** atoms, 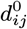 is the reference distance between these atoms, as found in the original structure, *γ* is the spring constant, and ***R_c_*** is a cutoff distance, typically between 3.5 Å and 15 Å. By default we let this value to 5 Å. However, if there are loosely connected structural fragments in the system, it makes sense to increase this value to 10 Å or even more. The Hessian matrix corresponding to this potential function is composed of the following blocks (21–23),

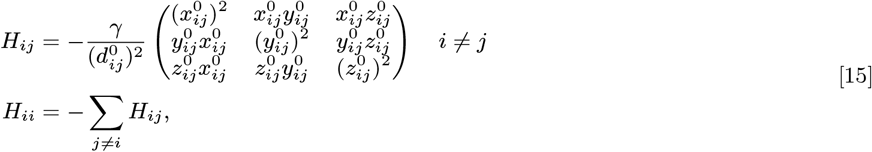

where *x_ij_* = *x_i_* − *x_j_, y_ij_* = *y_i_* − *y_j_*, and *z_ij_* = *z_i_* − *z_j_*. To rapidly compute this matrix, we use an efficient neighbor search algorithm (24).

### 3. Assessment of the transitions

#### A. Transition coverage

To assess the quality of the computed transitions, we measure the extent to which they cover the conformational deviation between the aligned starting and target states. Transition coverage is computed as

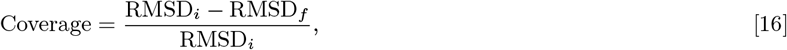

where RMSD_*i*_ is the initial root mean square deviation between the starting and target structures, and RMSD_*f*_ is the deviation between the final structure obtained from the computed transition and the target structure. The coverage varies between 0 (null prediction) and 1 (perfect prediction).

#### B. Collectivity

Collective motions can be characterised by their *collectivity **κ***, which is proportional to the exponential of the information entropy (25). The collectivity of a transition between two structures of a molecule with ***N*** atoms can be computed (26) as

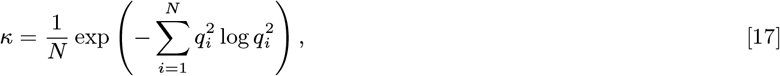

where ***q_i_*** are scaled Cartesian displacements of individual atoms, 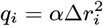, with the normalization factor ***α*** taken such that 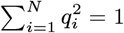. ***N_κ_*** gives an effective number of nonzero displacements 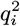. Thus, ***κ*** is confined to the interval {1/***N***;1}. If ***κ*** = 1, then the corresponding transition is maximally collective and has all the displacements 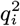 identical, which happens for rigid-body motions, for example. In the limit of an extremely localized motion, where only one single atom is affected, ***κ*** is minimal and equals to 1/***N***. In a similar way, one can estimate the degree of collectivity of a normal mode. For example, collectivity of the ***j***th mode is given by the same equation above provided that ***q_i_*** are now the scaled normal mode’s displacements,

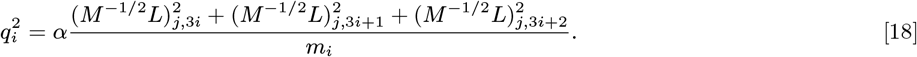

### 4. Computational details

For all the computations we used the NOLB package that rapidly performs linear and nonlinear NMA in Cartesian coordinates (15, 27). Given two states of a molecule in the PDB file format, we performed all the computations using the following commands, “NOLB initial.pdb final.pdb --linear” for the calculation of linear structural transitions, and “NOLB initial.pdb final.pdb --nlin” for the calculation of nonlinear structural transitions. Additional local minimization can be applied (with the “-m” flag) to keep the bond length and angles near the equilibrium positions. By default, structural transitions are computed between all the C***α*** atoms of the two structures, whose residues are aligned to each other using only sequence information. For a few cases from the test set 2, we identified ambiguities in the alignment leading to incorrect results. To resolve these cases, we performed an iterative alinement with 5 additional cycles, progressively excluding atoms with RMSD above a certain threshold (2 Å for 1he8:r, 2z0e:l and 1nw9:r, 4 Å for 1azs:r) at each cycle. The method is available free of charge for academic users on the three main platforms, MacOS, Linux, and Windows. We should also mention that the method is very rapid. For example, it took about 9 minutes to compute the nonlinear structural transitions for all proteins from the PPDBv5 (460 in total) with the local minimization enabled and using the 10 lowest-frequency normal modes on an Intel(R) Xeon(R) CPU E5-2630 v4 @ 2.20GHz. Performing 5 iterations of the multi-diagonalization scheme increased the computing time to about half an hour.

**Fig. S1.**
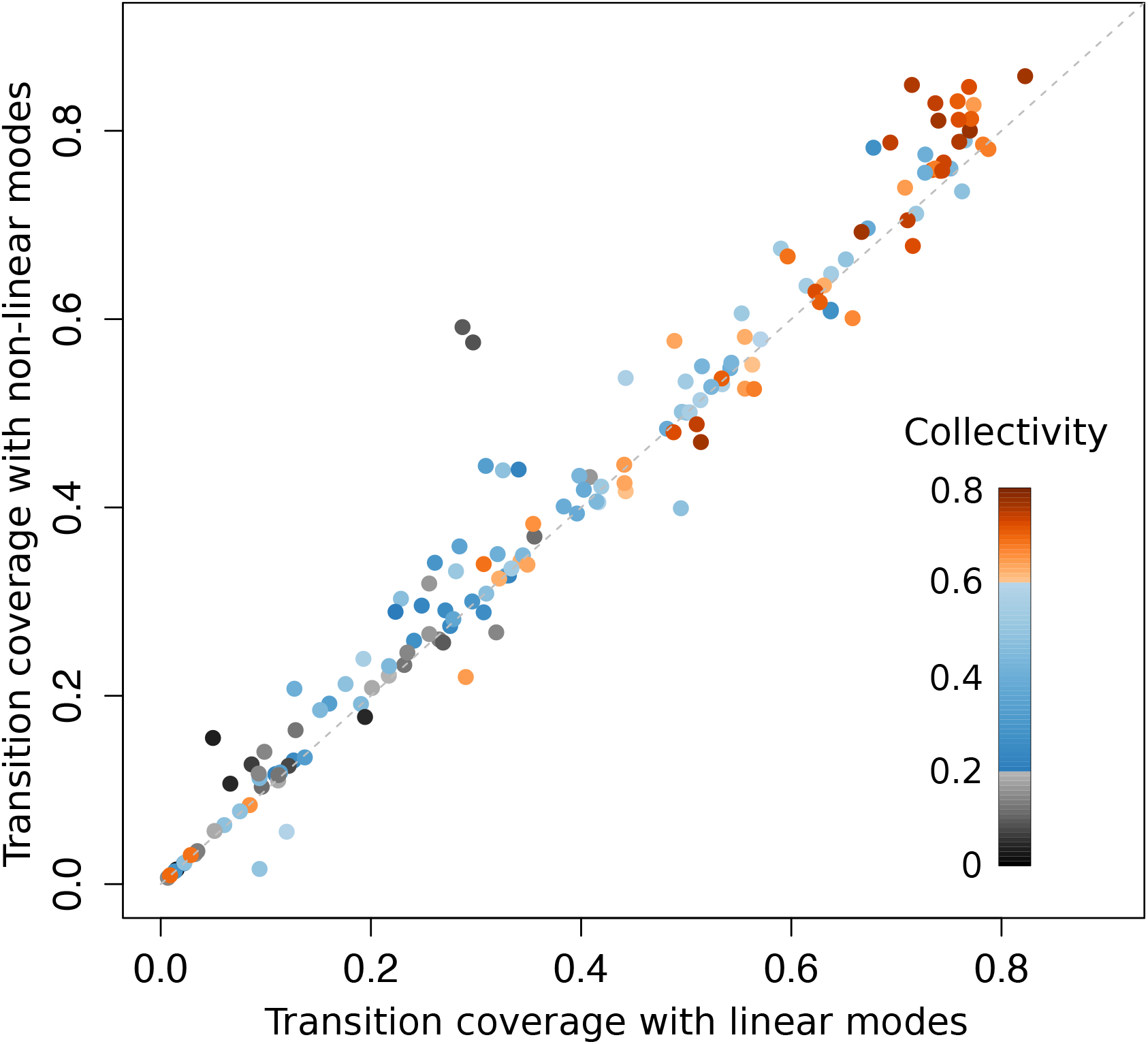
Comparison of transition coverage between the NOLB nonlinear modes and the classical linear modes. 155 structural transitions corresponding to 132 structure pairs were computed. The colors indicate the degrees of collectivity of the experimental transitions.

**Fig. S2.**
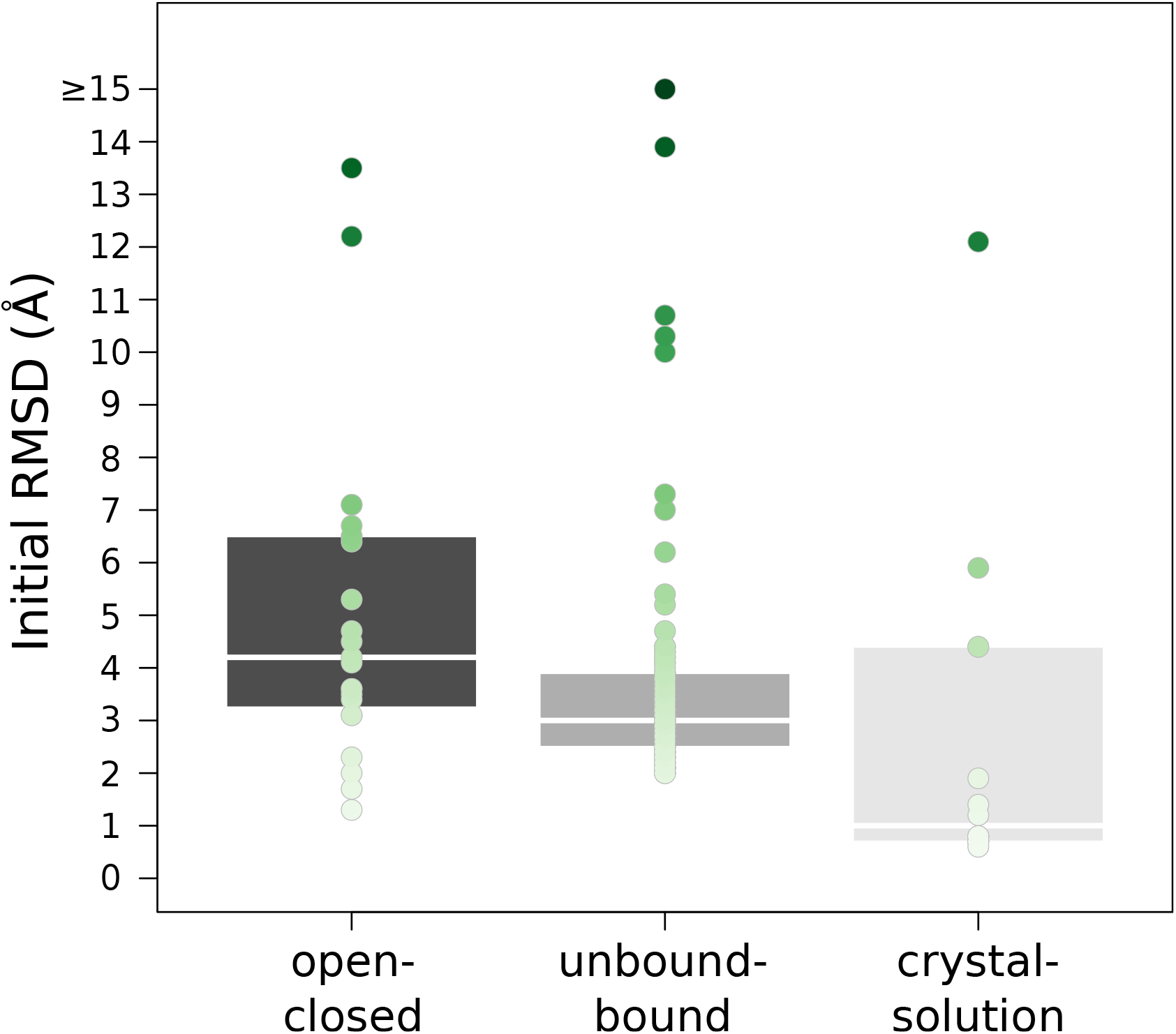
Distributions of initial RMSD between the starting and target structures in the three benchmark sets. Their detailed description of the benchmarks is given in *SI Appendix, Text 1*.

**Table S1.** Starting and target structures in the Cryo-EM 2015/2016 Model Challenge benchmark. The starting structures were solved by X-ray crystallography unless specified otherwise. The target structures result from the Model Challenge maps. For several targets, multiple starting templates were specified. For *β*-Galactosidase, two target cryo-EM structures were given in the Challenge. All the chains in the input structures were taken into account unless specified otherwise.

| Name | EMDB entry | Starting structure | Target structure |
| --- | --- | --- | --- |
| Tobacco Mosaic Virus | EMD-2842 | 1ei7:A<sup>1</sup> | 4udv |
| T20S Proteasome | EMD-5623 /<br>EMD-6287 | 1yar:A<sup>1</sup><br>1yar:H<sup>1</sup> | 3j9i:A<sup>1</sup><br>3j9i:H<sup>1</sup> |
| GroEL | EMD-6422 | 1ss8:A<sup>1</sup><br>1svt:A<sup>1</sup><br>1svt:H<sup>1</sup> | 3cau:A<sup>1</sup><br>3cau:A<sup>1</sup><br>3cau:A<sup>1</sup> |
| TRPV1 Channel | EMD-5778 | 6c8h<sup>2</sup> | 3j5p:A<sup>1</sup> |
| Brome Mosaic Virus | EMD-6000 | 1js9 | 3j7l |
| β-Galactosidase | EMD-5995 /<br>EMD-2984 | 1jz7:A<sup>1</sup><br>1jz7:A<sup>1</sup> | 3j7h:A<sup>1</sup><br>5a1a:A<sup>1</sup> |
| γ-Secretase | EMD-2677 /<br>EMD-3061 | 5fn2<sup>3</sup><br>5fn3<sup>3</sup><br>5fn4<sup>3</sup><br>5fn5<sup>3</sup> | 5a63<br>5a63<br>5a63<br>5a63 |

<sup>1</sup> In these structures, only a single chain was used. Its identifier is given after the colon.
<sup>2</sup> No starting X-ray template was specified in the Cryo-EM Model Challenge. Therefore, we used a homologous X-ray structure with about 50% of sequence identity. Two other similar structures can be retrieved from the PDB, 6c8f and 6c8g, but these are nearly identical to 6c8h.
<sup>3</sup> No homologous X-ray structures were found in the PDB. Thus, we used cryo-EM homologous structures.

**Movie S1. Nonlinear transition predicted for coagulation factor VIIa. The starting and target structures are colored in grey and black, respectively. The part of the transition in orange is obtained from the normal modes computed on the starting structure. The part in red is obtained by updating the modes three times. This transition was produced using the command “NOLB 1fak_r_u.pdb 1fak_r_b.pdb -n 10 --nlin --nIter 3 -m”.**

**Movie S2. Linear transition predicted for actin. The starting and target structures are colored in grey and black, respectively. The transition obtained with the classical linear modes is colored in blue. Please note the incorrect size of the loop that increases with the progression of the transition. This transition was produced using the command “NOLB 1atn_r_u.pdb 1atn_r_b.pdb -n 10 --linear --trajectory -s 33”.**

**Movie S3. Nonlinear transition predicted for actin. The starting and target structures are colored in grey and black, respectively. The transition obtained with NOLB nonlinear modes is colored in orange. This transition was produced using the command “NOLB 1atn_r_u.pdb 1atn_r_b.pdb -n 10 --nlin -m”.**

**Movie S4. Nonlinear transition predicted for diaminopimelate dehydrogenase. The starting and target structures are colored in grey and black, respectively. The part of the transition in orange is obtained from the normal modes computed on the starting structure. The part in red is obtained by updating the modes five times. This transition was produced using the command “NOLB 1dap.pdb 3dap.pdb -n 10 --nlin --niter 5 -m”.**

**Movie S5. Nonlinear transition predicted for the calcium ATPase pump. The residues undergoing the highest displacements are highlighted in pink color and stick representation. This transition was produced using the command “NOLB 1su4.pdb 1t5s.pdb -n 10 --nlin”.**

**Movie S6. Linear transition predicted for the calcium ATPase pump. The residues undergoing the highest displacements are highlighted in magenta color and stick representation. Please note unphysical scaling of the highlighted fragments with the progression of the transition. This transition was produced using the command “NOLB 1su4.pdb 1t5s.pdb -n 10 --linear --trajectory -s 60”.**

## References

1. Berman HM, et al. (2000) the Protein Data Bank. Nucleic Acids Res. 28:235–242.

2. Callaway E (2015) the Revolution Will Not Be Crystallized: A New Method Sweeps Through Structural Biology. Nature 525(7568):172–174.

3. Hayward S, Go N (1995) Collective variable description of native protein dynamics. Annual Review of Physical Chemistry 46(1):223–250.

4. Tama F, Miyashita O, III CLB (2004) Normal Mode Based Flexible Fitting of High-Resolution Structure into Low-Resolution Experimental Data from Cryo-EM. Journal of Structural Biology 147(3):315–326.

5. Tama F, Miyashita O, Brooks III CL (2004) Flexible multi-scale fitting of atomic structures into low-resolution electron density maps with elastic network normal mode analysis. Journal of Molecular Biology 337(4):985–999.

6. Suhre K, Navaza J, Sanejouand YH (2006) NORMA: a tool for flexible fitting of high-resolution protein structures into low-resolution electron-microscopy-derived density maps. Acta Crys-tallographica Section D: Biological Crystallography 62(9):1098–1100.

7. Schröder GF, Brunger AT, Levitt M (2007) Combining efficient conformational sampling with a deformable elastic network model facilitates structure refinement at low resolution. Structure 15(12):1630–1641.

8. Schröder GF, Levitt M, Brunger AT (2012) Super-resolution biomolecular crystallography with low-resolution data. Nature 464(7292):1218.

9. Tan RKZ, Devkota B, Harvey SC (2008) YUP. SCX: coaxing atomic models into medium resolution electron density maps. Journal of Structural Biology 163(2):163–174.

10. Zheng W (2011) Accurate flexible fitting of high-resolution protein structures into cryo-electron microscopy maps using coarse-grained pseudo-energy minimization. Biophysical Journal 100(2):478–488.

11. Lopéz-Blanco JR, Chacón P (2013) iMODFIT: efficient and robust flexible fitting based on vibrational analysis in internal coordinates. Journal of Structural Biology 184(2):261–270.

12. Gorba C, Miyashita O, Tama F (2008) Normal-mode flexible fitting of high-resolution structure of biological molecules toward one-dimensional low-resolution data. Biophysical Journal 94(5):1589–1599.

13. Kondrashov DA, Van Wynsberghe AW, Bannen RM, Cui Q, Phillips Jr GN (2007) Protein structural variation in computational models and crystallographic data. Structure 15(2):169–177.

14. Mendez R, Bastolla U (2010) Torsional network model: normal modes in torsion angle space better correlate with conformation changes in proteins. Physical Review Letters 104(22):228103.

15. Zhou L, Liu Q (2014) Aligning experimental and theoretical anisotropic b-factors: water models, normal-mode analysis methods, and metrics. The Journal of Physical Chemistry B 118(15):4069–4079.

16. Cavasotto CN, Kovacs JA, Abagyan RA (2005) Representing receptor flexibility in ligand docking through relevant normal modes. Journal of the American Chemical Society 127(26):9632–9640.

17. Mustard D, Ritchie DW (2005) Docking essential dynamics eigenstructures. Proteins Struct. Funct. Bioinf. 60(2):269–274.

18. Emekli U, Schneidman-Duhovny D, Wolfson H, Nussinov R, Haliloglu T (2008) HingeProt: Automated prediction of hinges in protein structures. Proteins Struct. Funct. Bioinf. 70(4):1219–1227.

19. Schneidman-Duhovny D, Nussinov R, Wolfson HJ (2007) Automatic prediction of protein interactions with large scale motion. Proteins Struct. Funct. Bioinf. 69(4):764–773.

20. Moal IH, Bates PA (2010) SwarmDock and the use of normal modes in protein-protein docking. International Journal of Molecular Sciences 11:3623–3648.

21. Fiorucci S, Zacharias M (2010) Binding site prediction and improved scoring during flexible protein–protein docking with ATTRACT. Proteins Struct. Funct. Bioinf. 78(15):3131–3139.

22. May A, Zacharias M (2008) Energy minimization in low-frequency normal modes to efficiently allow for global flexibility during systematic protein–protein docking. Proteins Struct. Funct. Bioinf. 70(3):794–809.

23. Neveu E, et al. (2018) RapidRMSD: Rapid determination of RMSDs corresponding to motions of flexible molecules. Bioinformatics 34(16):2757–2765.

24. Delarue M, Dumas P (2004) On the use of low-frequency normal modes to enforce collective movements in refining macromolecular structural models. Proceedings of the National Academy of Sciences 101(18):6957–6962.

25. Lindahl E, Azuara C, Koehl P, Delarue M (2006) NOMAD-Ref: Visualization, deformation and refinement of macromolecular structures based on all-atom normal mode analysis. Nucleic Acids Res. 34(2):W52–W56.

26. Lindahl E, Delarue M (2005) Refinement of docked protein–ligand and protein–dna structures using low frequency normal mode amplitude optimization. Nucleic Acids Res. 33(14):4496–4506.

27. Maschiach E, Nussinov R, Haim W (2010) Fiberdock: Flexible induced-fit backbone refinement in molecular docking. Proteins Struct. Funct. Bioinf. 78(6):1503–1519.

28. Venkatraman V, Ritchie DW (2012) Flexible protein docking refinement using pose-dependent normal mode analysis. Proteins Struct. Funct. Bioinf. 80(9):2262–2274.

29. Ma J (2005) Usefulness and limitations of normal mode analysis in modeling dynamics of biomolecular complexes. Structure 13(3):373–380.

30. Tama F, Sanejouand YH (2001) Conformational change of proteins arising from normal mode calculations. Protein Engineering 14(1):1–6.

31. Bonomi M, Pellarin R, Vendruscolo M (2018) Simultaneous Determination of Protein Structure and Dynamics Using Cryo-Electron Microscopy. Biophysical Journal 114(7):1604–1613.

32. Amadei A, Linssen AB, Berendsen HJ (1993) Essential dynamics of proteins. Proteins: Struct., Funct., Genet. 17(4):412–425.

33. Stepanova M (2007) Dynamics of essential collective motions in proteins: theory. Physical Review E 76(5 Pt 1):051918.

34. Spiwok V, Lipovova P, Kralova B (2007) Metadynamics in essential coordinates: free energy simulation of conformational changes. The Journal of Physical Chemistry B 111(12):3073–3076.

35. Fiorin G, Klein ML, Hénin J (2013) Using collective variables to drive molecular dynamics simulations. Molecular Physics 111(22-23):3345–3362.

36. Hoffmann A, Grudinin S (2017) NOLB: Nonlinear rigid block normal-mode analysis method. Journal of Chemical Theory and Computation 13(5):2123–2134.

37. López-Blanco JR, Aliaga JI, Quintana-Ortí ES, Chacón P (2014) iMODS: internal coordinates normal mode analysis server. Nucleic acids research 42(W1):W271–W276.

38. Frezza E, Lavery R (2015) Internal normal mode analysis (iNMA) applied to protein conformational flexibility. Journal of Chemical Theory and Computation 11(11):5503–5512.

39. Noguti T, Gõ N (1983) Dynamics of native globular proteins in terms of dihedral angles. Journal of the Physical Society of Japan 52(9):3283–3288.

40. Levitt M, Sander C, Stern PS (1985) Protein normal-mode dynamics: Trypsin inhibitor, cram-bin, ribonuclease and lysozyme. Journal of Molecular Biology 181(3):423–447.

41. Kamiya K, Sugawara Y, Umeyama H (2003) Algorithm for normal mode analysis with general internal coordinates. Journal of Computational Chemistry 24(7):826–841.

42. Lu M, Poon B, Ma J (2006) A new method for coarse-grained elastic normal-mode analysis. Journal of Chemical Theory and Computation 2(3):464–471.

43. Lopéz-Blanco JR, Garzón JI, Chacón P (2011) iMod: Multipurpose normal mode analysis in internal coordinates. Bioinformatics 27(20):2843–2850.

44. Vreven T, et al. (2015) Updates to the integrated protein-protein interaction benchmarks: Docking benchmark version 5 and affinity benchmark version 2. Journal of Molecular Biology 427(19):3031–3041.

45. Lawson C, et al. (2018) CryoEM Models and Associated Data Submitted to the 2015/2016 EMDataBank Model Challenge. Zenodo. https://doi.org/10.5281/zenodo.1165999.

46. Tirion MM (1996) Large amplitude elastic motions in proteins from a single-parameter, atomic analysis. Physical Review Letters 77(9):1905.

47. Krebs WG, et al. (2002) Normal mode analysis of macromolecular motions in a database framework: developing mode concentration as a useful classifying statistic. Proteins 48(4):682–695.

48. Durand P, Trinquier G, Sanejouand YH (1994) A new approach for determining low-frequency normal modes in macromolecules. Biopolymers 34(6):759–771.

49. Tama F, Gadea FX, Marques O, Sanejouand YH (2000) Building-block approach for determining low-frequency normal modes of macromolecules. Proteins: Structure, Function and Bioinformatics 41(1):1–7.

50. Neveu E, Ritchie DW, Popov P, Grudinin S (2016) PEPSI-Dock: A detailed data-driven protein-protein interaction potential accelerated by polar fourier correlation. Bioinformatics 32(17):i693–i701.

51. Dobbins SE, Lesk VI, Sternberg MJE (2008) Insights into protein flexibility: the relationship between normal modes and conformational change upon protein–protein docking. Proceedings ofthe National Academy of Sciences of the United States of America 105(30):10390–10395.

52. Marsh JA, Teichmann SA, Forman-Kay JD (2012) Probing the diverse landscape of protein flexibility and binding. Current Opinion in Structural Biology 22(5):643–650.

53. Yang L, Song G, Jernigan RL (2007) How well can we understand large-scale protein motions using normal modes of elastic network models? Biophysical Journal 93(3):920–929.

## References

1. Lopéz-Blanco JR, Garzón JI, Chacón P (2011) iMod: Multipurpose normal mode analysis in internal coordinates. Bioinformatics 27(20):2843–2850.

2. Tekpinar M (2018) Flexible fitting to cryo-electron microscopy maps with coarse-grained elastic network models. Molecular Simulation pp. 1–9.

3. Echols N, Milburn D, Gerstein M (2003) MolMovDB: Analysis and visualization of conformational change and structural flexibility. Nucleic Acids Research 31(1):478–482.

4. Chen VB, et al. (2010) Molprobity: All-atom structure validation for macromolecular crystallography. Acta Crystallographica Section D Structural Biology 66(1):12–21.

5. Vreven T, et al. (2015) Updates to the integrated protein-protein interaction benchmarks: Docking benchmark version 5 and affinity benchmark version 2. Journal of Molecular Biology 427(19):3031–3041.

6. Dobbins SE, Lesk VI, Sternberg MJE (2008) Insights into protein flexibility: the relationship between normal modes and conformational change upon protein–protein docking. Proceedings of the National Academy of Sciences of the United States of America 105(30):10390–10395.

7. Lawson C, et al. (2018) CryoEM Models and Associated Data Submitted to the 2015/2016 EMDataBank Model Challenge. Zenodo. https://doi.org/10.5281/zenodo.1165999.

8. Noguti T, Gõ N (1983) Dynamics of native globular proteins in terms of dihedral angles. Journal of the Physical Society of Japan 52(9):3283–3288.

9. Levitt M, Sander C, Stern PS (1985) Protein normal-mode dynamics: Trypsin inhibitor, crambin, ribonuclease and lysozyme. Journal of Molecular Biology 181(3):423–447.

10. Lu M, Poon B, Ma J (2006) A new method for coarse-grained elastic normal-mode analysis. Journal of Chemical Theory and Computation 2(3):464–471.

11. Mendez R, Bastolla U (2010) Torsional network model: normal modes in torsion angle space better correlate with conformation changes in proteins. Physical Review Letters 104(22):228103.

12. Durand P, Trinquier G, Sanejouand YH (1994) A new approach for determining low-frequency normal modes in macromolecules. Biopolymers 34(6):759–771.

13. Tama F, Gadea FX, Marques O, Sanejouand YH (2000) Building-block approach for determining low-frequency normal modes of macromolecules. Proteins: Structure, Function and Bioinformatics 41(1):1–7.

14. Li G, Cui Q (2002) A coarse-grained normal mode approach for macromolecules: An efficient implementation and application Ca^2^+-ATPase. Biophysical Journal 83(5):2457–2474.

15. Hoffmann A, Grudinin S (2017) NOLB: Nonlinear rigid block normal-mode analysis method. Journal of Chemical Theory and Computation 13(5):2123–2134.

16. Lopéz-Blanco JR, Chacón P (2013) iMODFIT: efficient and robust flexible fitting based on vibrational analysis in internal coordinates. Journal of Structural Biology 184(2):261–270.

17. López-Blanco JR, Aliaga JI, Quintana-Ortí ES, Chacón P (2014) iMODS: internal coordinates normal mode analysis server. Nucleic acids research 42(W1):W271–W276.

18. Song G, Jernigan RL (2006) An enhanced elastic network model to represent the motions of domain-swapped proteins. Proteins: Structure, Function, and Bioinformatics 63(1):197–209.

19. Yang L, Song G, Jernigan RL (2007) How well can we understand large-scale protein motions using normal modes of elastic network models? Biophysical Journal 93(3):920–929.

20. Zheng W (2011) Accurate flexible fitting of high-resolution protein structures into cryo-electron microscopy maps using coarse-grained pseudo-energy minimization. Biophysical Journal 100(2):478–488.

21. Doruker P, Atilgan AR, Bahar I (2000) Dynamics of proteins predicted by molecular dynamics simulations and analytical approaches: Application to *α*-amylase inhibitor. Proteins: Structure, Function and Bioinformatics 40(3):512–524.

22. Atilgan AR, et al. (2001) Anisotropy of fluctuation dynamics of proteins with an elastic network model. Biophysical Journal 80(1):505–515.

23. Bahar I, Lezon TR, Bakan A, Shrivastava IH (2010) Normal mode analysis of biomolecular structures: Functional mechanisms of membrane proteins. Chemical Reviews 110(3):1463–1497.

24. Artemova S, Grudinin S, Redon S (2011) A comparison of neighbor search algorithms for large rigid molecules. Journal of Computational Chemistry 32(13):2865–2877.

25. Brüschweiler R (1995) Collective protein dynamics and nuclear spin relaxation. The Journal of Chemical Physics 102(8):3396–3403.

26. Tama F, Sanejouand YH (2001) Conformational change of proteins arising from normal mode calculations. Protein Engineering 14(1):1–6.

27. Neveu E, et al. (2018) RapidRMSD: Rapid determination of RMSDs corresponding to motions of flexible molecules. Bioinformatics 34(16):2757–2765.

